# Disrupted social memory ensembles in the ventral hippocampus underlie social amnesia in autism-associated *Shank3* mutant mice

**DOI:** 10.1101/2021.06.25.449869

**Authors:** Kentaro Tao, Myung Chung, Akiyuki Watarai, Ziyan Huang, Mu-Yun Wang, Teruhiro Okuyama

## Abstract

The ability to remember conspecifics is critical for adaptive cognitive functioning and social communication, and impairments of this ability are hallmarks of autism spectrum disorders (ASDs). Although hippocampal ventral CA1 (vCA1) neurons are known to store social memories, how their activities are coordinated remains unclear. Here we show that vCA1 social memory neurons, characterized by enhanced activity in response to memorized individuals, were preferentially reactivated during sharp-wave ripples (SPW-Rs). Spike sequences of these social replays reflected the temporal orders of neuronal activities within theta cycles during social experiences. In ASD model *Shank3* knockout mice, the proportion of social memory neurons was reduced, and neuronal ensemble spike sequences during SPW-Rs were disrupted, which correlated with impaired discriminatory social behavior. These results suggest that SPW-R-mediated sequential reactivation of neuronal ensembles is a canonical mechanism for coordinating hippocampus-dependent social memories and its disruption underlies the pathophysiology of social memory defects associated with ASD.

## INTRODUCTION

The ability to recognize and memorize familiar conspecifics is crucial for animals that engage in social interactions [1, 2]. Several brain regions have been identified as being associated with the experience-dependent encoding of social characteristics in rodents [3, 4]. In particular, the ventral CA1 sub-region of the hippocampus (vCA1) contains neurons that respond to other individuals [5] but not to inanimate objects [6] and has been identified as a key locus for social memory storage [5, 7]. *In vivo* Ca^2+^ imaging in freely moving mice revealed that a subset of vCA1 neurons is activated by the presence of familiar mice during social interactions [5]; however, how the activities of these social neurons are organized and maintained at the fine temporal resolution necessary for effective memory encoding and recall remains unknown.

Hippocampal sharp-wave ripples (SPW-Rs) are transient, high-frequency, field oscillations that are typically observed during slow-wave sleep and quiet wakefulness [8] and play a pivotal role in the formation of episodic memories encoded in the hippocampus. In the dorsal hippocampus, place-cell ensembles that are activated during awake exploration are sequentially reactivated during SPW-Rs that occur during both slow-wave sleep [9–11] and transient immobility during awake exploration [12, 13]. SPW-Rs have been suggested to play prominent roles in synaptic plasticity [14] and memory consolidation [15, 16]; they can also be observed in the ventral hippocampus [17], where they are thought to distribute behavior-contingent information to multiple downstream regions [18]. Hippocampal network oscillations, such as theta waves, can propagate along the dorsoventral axis of the hippocampus [19]; however, the SPW-Rs that arise in the ventral segment of the hippocampus often remain isolated [17]. To date, it remains unclear whether the ventral hippocampus neuronal activity during SPW-Rs inherently represents spatial information or is used for processing related to other aspects, such as social information.

Here, we report that vCA1 neurons that respond to familiar social targets are preferentially co-activated during SPW-Rs following social memory formation. The relative spike timing of these social neurons was preserved across SPW-Rs, indicating that social information is likely organized based on a temporal code. Furthermore, in SH3 and multiple ankyrin repeat domains 3 (*Shank3*)-knockout (KO) mice, a genetic model of autism spectrum disorder (ASD), a number of social neurons were reduced, and vCA1 neurons exhibited a loss of coactivation fidelity during SPW-Rs that was correlated with impaired social discrimination behavior. These results indicate that social information is processed in the ventral hippocampus through a mechanism that is analogous to that used to process spatial information in the dorsal hippocampus and that this mechanism for social information processing is impaired in a genetic model of ASD.

## MATERIALS AND METHODS

### Mice

All procedures were in accordance with protocols approved by the Institutional Animal Care and Use Committee at the Institute for Quantitative Biosciences, University of Tokyo. All animals were socially housed under 12 h (7 a.m.–7 p.m.) light/dark cycle conditions, with ad libitum access to food and water. C57BL/6J wild-type mice were obtained from the Central Laboratories for Experimental Animals, Japan, and *Shank3*-KO mice were obtained from the Jackson laboratory (B6.129-Shank3^tm2Gfng^/J, Stock No: 017688). The *Shank3*-KO allele has a neomycin cassette replacing the PDZ domain (exons 13–16) of the *Shank3* gene. Five adult male C57BL/6J mice (28–32 g, 3–4 months old) and four adult male homozygous *Shank3*-KO mice (26–32 g, 3–4 months old) were used as subject mice. Young adult male C3H and BALB/c mice (22–28 g, 6–8 weeks old) were used as stimulator mice.

### Surgical procedures

All surgical procedures were performed under 1–2% isoflurane anesthesia and with the mice mounted on a stereotaxic apparatus (SR-9M-HT; Narishige, Tokyo, Japan). Dental cement was used to attach a custom-made head frame onto a cleared skull for the convenience of subsequent silicon probe implantation. Mice were individually housed after the surgery. After 1–2 weeks of recovery, mice were again mounted with the head frame, and 64-channel high-density silicon probes (A4×16-Poly2-5mm-20s-150-160; NeuroNexus, Ann Arbor, MI) targeting the right ventral hippocampus were implanted parallel to the midline (AP: −3.00–3.45 mm, ML: +3.10 mm from bregma). Probes were attached to movable microdrives and lowered gradually over the course of several days until the CA1 pyramidal layer was reached, determined by the appearance of hippocampal SPW-Rs and pyramidal cell activity.

### Social discrimination test (SDT)

Mice were individually habituated to the investigator by handling for several minutes on each of two consecutive days (Day-1 and Day-2). Habituation to the social arena and recording cable was performed for 5 min on each of two consecutive days (Day-2 and Day-3). The social arena was a white acrylic box (area, 40×40 cm^2^, height, 30 cm); two custom-made social chambers with quadrant-shaped bottoms (7.5 cm radius) printed with a 3D printer (Original Prusa i3 MK3S; Prusa Research Prague, Czech) were placed at the opposite corners of the arena. On Day-4, a stimulator mouse (C3H; A) was placed into the home cage of the subject mouse for familiarization for 2 h. Thereafter, the familiarized mouse (A) and a novel mouse (BALB/c; B) were placed in the left or right chamber, and the subject mouse was allowed to explore inside the arena for 5 min (A–A trial). This procedure was repeated by alternating the position of the two stimulator mice (B–A trial) and then in their absence (E–E trial). Trials were recorded at 25 frames per second from the top view of the arena using Bonsai [20]. Body positions of subject mice were detected using DeepLabCut [21].

### *In vivo* electrophysiology

During the SDT and the subsequent resting period, mice were connected to a 64-channel amplifier board (RHD2164; Intan Technologies, Los Angeles, CA), and neural activities were sampled at 30 kHz using the Open Ephys data acquisition system [22]. The wide-band signal was downsampled to 1.25 kHz and used as the LFP signal. Spike sorting was performed automatically using Kilosort2 [23]. This was followed by manual adjustment of the waveform clusters using the phy graphical user interface [24]. Units with firing rate less than 15 Hz and trough-to-peak length of spike shapes longer than 0.5 ms were classified as excitatory neurons [25, 26].

### Histology

After completion of the experiments, mice were deeply anesthetized and transcardially perfused with phosphate-buffered saline and then 4% paraformaldehyde. Thereafter, brains were dissected out and post-fixed in 4% paraformaldehyde at 4 °C overnight, and fixed samples were sectioned into 100-μm coronal slices using a vibratome (Leica, VT1000S; Leica Microsystems, Wetzlar, Germany). For verifying the electrode tracks, images were acquired using a microscope (BZ-X710; Keyence, Tokyo, Japan).

### Behavioral analysis

All analyses were performed using MATLAB unless otherwise stated. The relative distance (*d*) between the subject mouse and two stimulator mice in each frame was calculated as (*d_A_* — *d_B_*)/(*d_A_* + *d_B_*), where *d_A_* and *d_B_* are the distances between the body position of the subject and the corners of social arena where stimulator mouse A or B, respectively, were placed. Coordinates with |d| ≥ 0.5 (corresponding approximately to twice the social chamber’s radius) were defined as being in the social zone. Social preference of the subject mouse was computed as (*Dur_A_* — *Dur_B_*)/(*Dur_A_* + *Dur_B_*), where *Dur_A_* and *Dur_B_* are the total time spent in the social zone around the stimulator mouse A or B, respectively.

The subject’s egocentric direction relative to each stimulator mouse was calculated by the relative orientation of the tail-to-body vector of the subject toward the corners of arena where stimulator mice were placed. Nose positions of the subject that were occasionally occluded by the head-mounted apparatus and made detection likelihood unstable were not utilized for the analysis.

### Social activity of hippocampal units

Neurons having mean firing rates lower than 15 Hz and trough-to-peak spike shape lengths longer than 0.5 ms were considered to be pyramidal neurons. Social preference score for each unit was computed as (*f_A_* — *f_B_*)/(*f_A_* + *f_B_*), where *f_A_* and *f_B_* were the firing rates of a unit when the subject was located in the social zone around the stimulator mouse A or B, respectively. Significance of the calculated preference was determined by a permutation test of spike counts in each video frame (repeated 10,000 times). The P value represents the proportion of shuffled values larger than the actual observed value.

The egocentric directional tuning function for each unit was the ratio between the peaks of spike count and total time spent in each direction in bins of 5° and smoothed with a Gaussian kernel of 2 bins.

### LFP spectrum analysis

Root mean square (RMS) of the LFP signal across channels having pyramidal cells was calculated at each time point. The signal was then binned into 10-s epochs and the instantaneous power spectrum density (PSD) was estimated using Welch’s method with a Hamming window (size = 1.33 s, overlap = 0.67 s). The obtained PSD was used to display spectrograms across offline recordings, to compare the spectral structure of the LFP between genotypes, and to assess the behavioral state during offline recordings as described below. To display spectrograms of SPW-Rs, continuous wavelet transform (CWT) was applied to unfiltered LFP signals using complex Morlet wavelets with a parameter of 7.

### Sleep analysis

To assess the behavioral state during offline recordings, signals from a 3-axis accelerometer on the headstage were utilized to calculate the derivatives of roll and pitch, which reflect angular velocities of the subject’s head. The L2 norm of these derivatives was used as a proxy for the overall movement of the subject. The signal was then binned into 10-s epochs and sleep epochs were defined as the period of sustained immobility (0.1 arbitrary unit/s). The delta (2–5 Hz) power calculated by Welch’s method was referenced to confirm that the movement signal recapitulated the physiological sleeping state.

### Ripple event detection

The LFP signal was bandpass filtered (150–250 Hz) on each channel having pyramidal neurons, and the root mean square across channels was calculated at each time point. The power was then Gaussian smoothed (4 ms standard deviation [s.d.]). Epochs with a peak power exceeding the mean by at least 5 s.d. for at least 15 ms were detected. The sample points at which the power reduced below 1 standard deviation were determined as the points of onset and offset of the epochs. Ripple pairs with peaks closer than 50 ms were merged into single events.

### Unit coactivation analysis

To measure the pairwise coactivation of units during online recordings, we used the cosine similarity between two vectors of spike counts, calculated as:

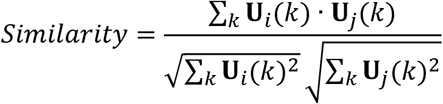

where ***U**_i_*(*k*) and ***U**_j_*(*k*) are spike counts of unit *U_i_*, and *U_j_* within the *k^th^* movie frame. Likewise, the pairwise coactivation of units during ripple events was evaluated using the same formula but with ***U**_i_*(*k*) and ***U**_j_*(*k*) being the spike counts of unit *U_i_*, and *U_j_* during the *k^th^* ripple event. For offline recordings, the significance of the calculated similarity was determined by a permutation test of spike counts during each ripple event (repeated 100,000 times). The P value represents the proportion of shuffled values larger than the actual observed value.

### Spike sequence analysis

The LFP signal was bandpass filtered (6–12 Hz) and averaged across channels having pyramidal cells. Theta cycles started at the peak of the LFP. Cycles shorter than 80 ms or longer than 200 ms were discarded, and only periods containing at least three contiguous cycles with both troughs and peaks larger than 1 s.d. were selected for further analysis. Awake sharp waves were excluded from the analysis by detecting ripples using the same criteria for offline recordings but with no constraint by MUAs. Rank order of a neuron in each online theta cycle or offline ripple event was defined as the normalized temporal position of a first spike in the sequence of all cells that participated in that theta cycle or ripple event. To calculate the difference of rank orders between cell pairs, the orders of the cells were determined so that the differences during ripple events were always a positive value.

## RESULTS

### Single-unit recording of vCA1 neurons during a social discrimination test

Mice display a natural inclination to approach a novel mouse rather than a familiar mouse. After 2 h of familiarization between the subject and a stimulator mouse in a home cage, we employed a social discrimination test (SDT) to assess the responses to the familiar mouse-A and a novel mouse-B (Fig. 1a). The SDT consisted of two consecutive 5 min trials in which the subject mouse was placed with mouse-A and mouse-B in counter-balanced positions (“social trials”) in a social arena, followed by a 5 min “control trial” with two empty social chambers. We confirmed that the subject mouse interacted with the novel mouse-B for a significantly longer period of time than with the familiar mouse-A (time spent in the social zone [mean ± SEM]: 145 ± 11 sec, mouse-A versus 185 ± 18 sec, mouse-B; *p* = 0.045, paired *t*-test; discrimination index: *p* = 0.025, Wilcoxon signed-rank test) (Fig. 1b-d), consistent with the findings of our previous study [5]. To analyze social memory representation at a fine temporal resolution, we implanted high-density, 64-site, four-shank silicon probes in the vCA1 (Fig. 1e) and recorded neural activity during the social memory task (“online” recording) as well as during the subsequent 2 h memory consolidation phase (“offline” recording). For each cell, we calculated a social preference score based on the body position of the subject mouse during the SDT; cells that exhibited significant activation in response to social interactions either with the familiar mouse-A (e.g., Unit #1 in Fig. 1f,g) or the novel mouse-B (e.g., Unit #7 in Fig. 1f,g) were termed “mouse-A cells” and “mouse-B cells”, respectively. Additionally, each cell was classified either as an excitatory pyramidal neuron or an inhibitory interneuron based on its firing rate and trough-to-peak length of spike shapes (Supplementary Fig. 1a). Of the 104 units recorded over 11 sessions, 75 units were classified as putative excitatory pyramidal neurons, whereas 29 units were classified as putative inhibitory interneurons. Among the putative pyramidal neurons, we identified 31.1 % as mouse-A cells and 4.0 % as mouse-B cells, and found that their overall activities indicated a significant preference for the familiar mouse-A (Wilcoxon signed-rank test, *p* = 4.9×10^-4^) (Fig. 1h), similar to our previous findings based on Ca^2+^ imaging [5]. In contrast, the familiar mouse-A and novel mouse-B cell fractions among the putative inhibitory interneurons did not differ significantly (mouse-A cells, 14.3 %; mouse-B cells 7.1 %; Wilcoxon signed-rank test, *p* = 0.66; Supplementary Fig. 1b).

**Fig. 1.**
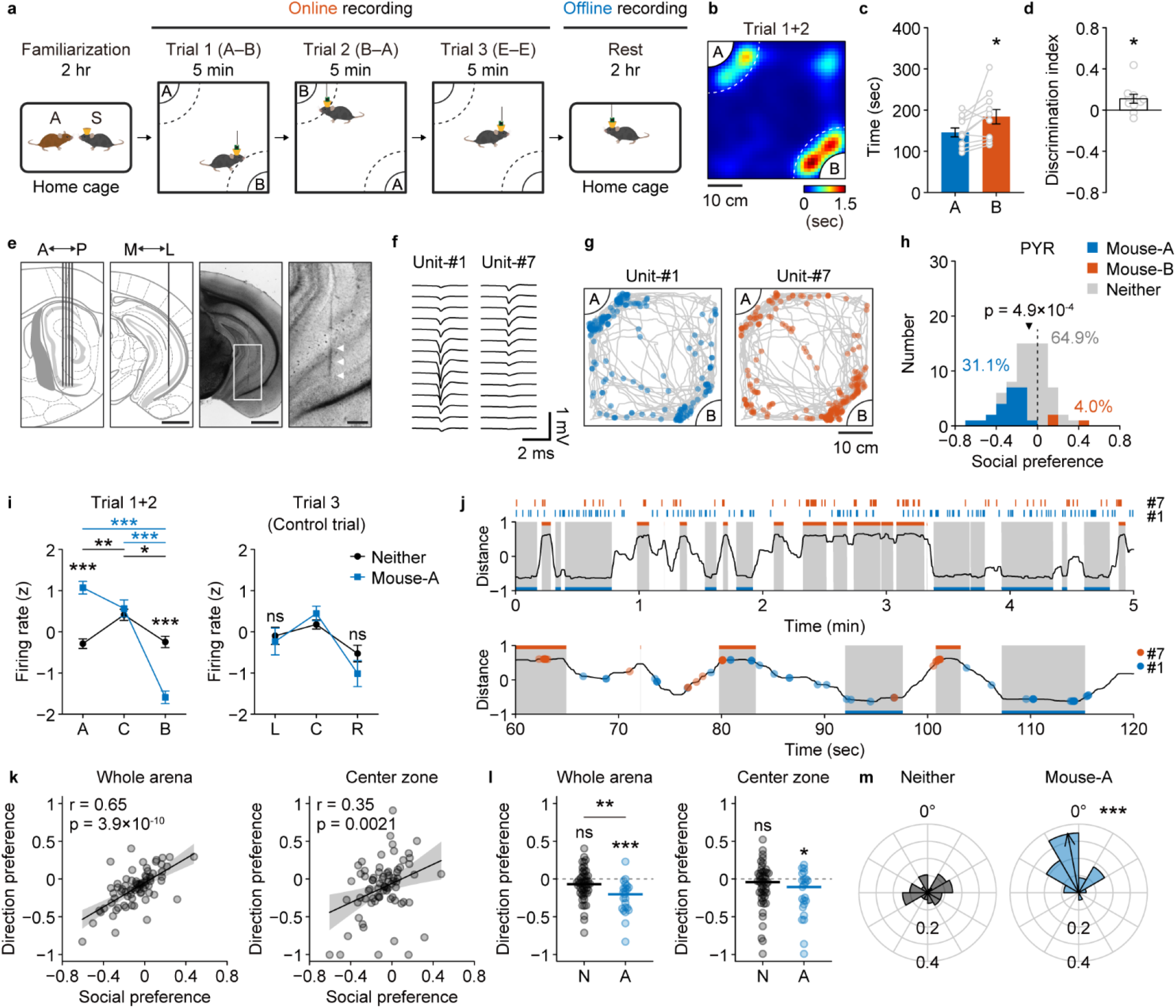
Electrophysiological recordings of ventral CA1 social neurons. **a** Schematic of the behavioral paradigm. After 2 h of familiarization between a subject mouse (S) and a stimulator mouse-A in a home cage, the subject mouse was allowed to explore a social arena where the familiar mouse-A and a novel mouse-B were placed. Dotted lines in the arena indicate the borders of social zones. **b** Heat map of occupancy time during Trial 1 and Trial 2 from a representative session. Note that the map for Trial 2 is rotated and superposed onto that of Trial 1. **c**, **d** Comparison of duration spent in the social zone (**c**) and the discrimination index (**d**). *n* = 11 sessions with 5 subjects; \**p* = 0.045, paired *t*-test (**c**) and \**p* = 0.025, Wilcoxon signed-rank test (**d**). Data are represented as mean ± SEM. **e** A four-shank silicon probe was implanted in the ventral hippocampus of freely moving mice. Arrowheads in the magnified view indicate the probe track. Scale bars: 1 mm (unmagnified images) and 200 μm (magnified images). **f** Representative recorded spike waveforms. **g** Spatial firing patterns units shown in **f**. Unit-#1, a mouse-A cell. Unit-#7, a mouse-B cell. Gray lines represent the trajectories of the subject mouse during Trials 1 and 2. **h** Social preference scores of recorded pyramidal neurons (n = 75 cells). Blue, mouse-A cells (*n* = 23 cells, 31.1 %); red, mouse-B cells (*n* = 3 cells, 4.0 %); grey, neither neurons (*n* = 48 cells, 64.9 %). The p value indicates that the median value of all pyramidal neurons was significantly different from zero (Wilcoxon signed-rank test). **i** Z-scored firing rates of mouse-A cells and neither cells during social trials (Trial 1+2; left) and the control trial (Trial 3; right). Trial 1+2: UnitType×Area, *F*_(1,69)_ = 96.6, *p* = 9.5×10^-15^; \*\*\**p* = 3.3×10^-9^ (A), 9.3×10^-8^ (B). Mouse-A: \*\*\**p* = 9.6×10^-10^ (A vs B), 2.1×10^-7^ (Center vs B); Neither: \*\**p* = 0.0081 (A vs Center), \**p* = 0.024 (Center vs B). Trial 3: UnitType×Area, F_(1,69)_ = 0.42, *p* = 0.52. All statistical tests are two-way repeated measures ANOVA and Tukey–Kramer multiple comparison tests. Data are represented as mean ± SEM. **j** Firing timings of the two units shown in **f**, and relative social distance of the subject mouse. Shaded periods indicate that the subject was in the social zone around mouse-A (blue) or mouse-B (red). **k** Correlations between social preferences and directional social preferences of pyramidal neurons (*n* = 75 cells). Whole arena, *r* = 0.65, *p* = 3.9×10^-10^; center zone, *r* = 0.35, *p* = 0.0021. **l** Comparison of directional social preferences. Comparison between neither vs mouse-A cells: \*\**p* =0.0022, Wilcoxon rank sum test. Comparison between each group vs zero: mouse-A cells in whole arena, \*\*\**p* =3.7×10^-4^ and mouse-A cells in center zone, \**p* = 0.026, Wilcoxon signed-rank test. ns, not significant. **m** Preferred directions and mean resultant vectors of neither cells and mouse-A cells. The black arrow direction indicates mean direction preference of all neither cells and mouse-A cells. Arrow lengths of mean resultant vectors indicate the direction concentration parameter, kappa. Neither: *n* = 48 cells, mean direction = 317.0°, *p* = 0.31, V test for circular uniformity against 0°; mouse-A: *n* = 23 cells, mean direction = 10.6°, \*\*\**p* = 1.3×10^-6^, V test for circular uniformity against 0°.

### Firing properties of vCA1 neurons during online social experiences

We then carefully examined the in vivo firing properties of vCA1 neurons during the SDT. Cells classified neither as mouse-A cells nor as mouse-B cells (“neither cells”) were equally active when the subject mouse approached either mouse-A or mouse-B. In contrast, mouse-A cells were more active when the subject mouse approached mouse-A and, conversely, were less active when subject mouse approached mouse-B (Fig. 1i and Supplementary Fig. 1). When the subject mouse explored the arena in the absence of any social targets (Trial 3), the activity of the mouse-A cells was similar to that of the neither cells. These data suggest that a fraction of vCA1 cells is adaptively modulated to be activated or inactivated by the presence of social cues, which gives rise to the specific responses to a memorized social target.

Closer investigation revealed that social cells fired both when the subject mouse was interacting with the preferred social target and when approaching it (Fig. 1j), which led us to examine the neuronal activity patterns in terms of the relative direction of the subject towards either social target in the arena. By calculating the directional social preference score of each cell, for which firing was considered only when the subject was oriented towards the social targets, we observed a significant positive correlation between the social preference scores and the directional social preference scores (Fig. 1k, left). This correlation was maintained even when we narrowed down the criteria to include activities recorded when subjects were in the center zone of the arena (Fig 1k, right), indicating that vCA1 neuronal activity related to social experiences appeared to be modulated not only by close interaction but also by distant recognition of social targets. Indeed, familiar mouse-A cells, determined based on social preference scores, showed a significant directional social preference as a population toward mouse-A throughout the arena (−0.21 ± 0.05, *p* = 0.0024, Wilcoxon’s signed-rank test) as well as in the center zone (−0.18 ± 0.07, *p* = 0.026, Wilcoxon’s signed-rank test) (Fig. 1l). Directional preferences relative to the social target also confirmed that the mouse-A cells preferentially fired when the subject was oriented towards mouse-A (Fig. 1m).

### Firing patterns of vCA1 neurons during offline SPW-R events

To determine how the activities of vCA1 social memory neurons (Fig. 2a) are coordinated and organized during SPW-R events, we investigated the firing patterns of pyramidal cells during offline recordings (Fig. 2b). In addition to the expected increase in delta power (1–4 Hz), which indicated that the subject was resting, a concurrent increase in ripple power (150–250 Hz) was observed (Fig. 2b and Supplementary Fig. 2). Across all sessions, we consistently observed that SPW-R events were accompanied by multi-unit activities during the offline recordings (Fig. 2c; 1.1 ± 0.5×10^3^ events per 2 h, *n* = 11 sessions). The familiar mouse-A cells showed a significantly higher firing rate around ripple peaks (Fig. 2d,e) and participated in a larger fraction of ripples (Fig. 2f) compared to neither cells, indicating that cells encoding social representations are preferentially reactivated following social experiences.

**Fig. 2.**
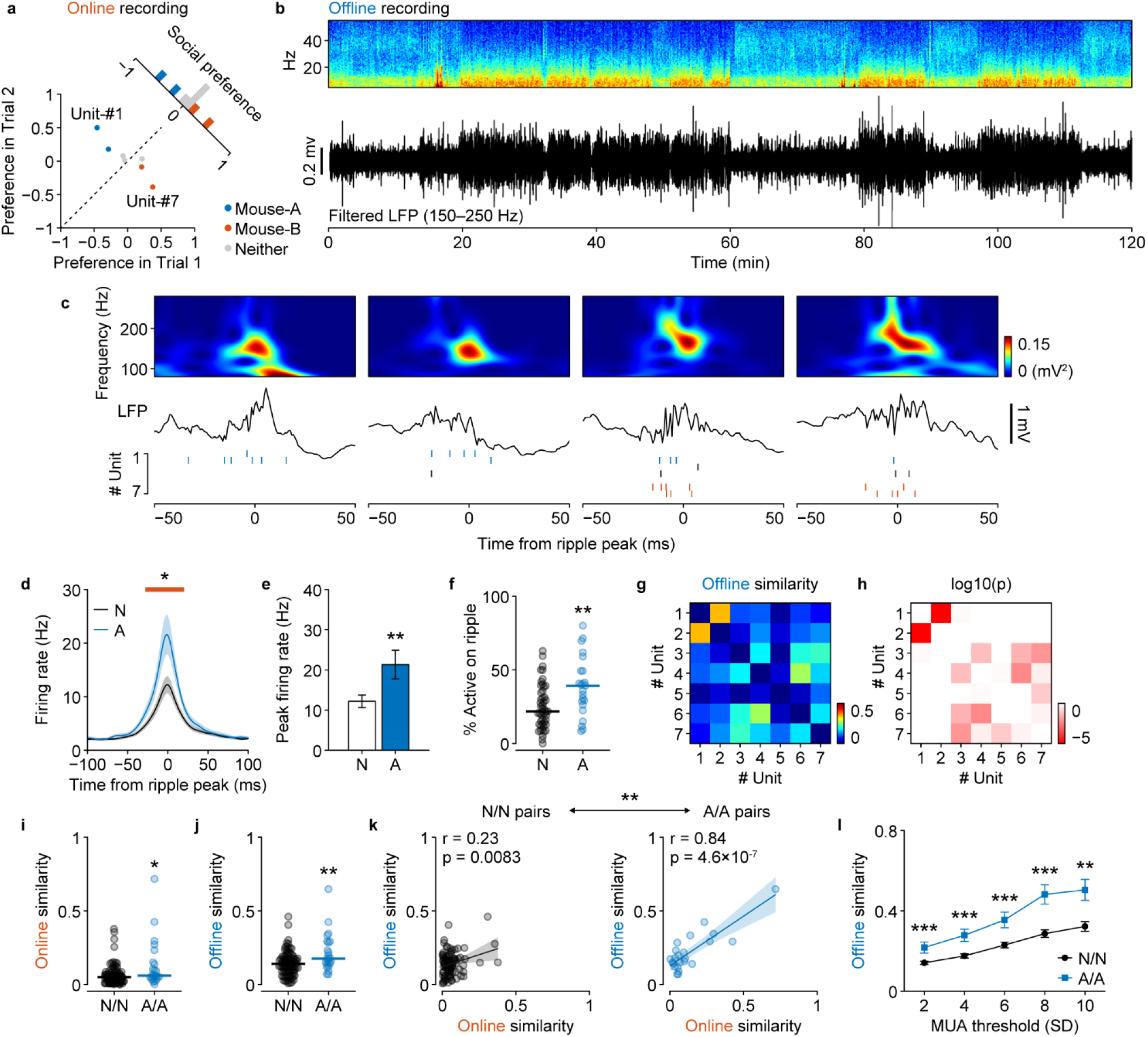
Neurons with similar social representations are co-activated during offline SPW-Rs. **a** Social preferences of example neurons recorded in the same session. **b** Example power spectrum and filtered ripple band trace of the LFP recorded in the home cage. **c** Representative SPW-Rs and raster plots of the units recorded during the same session as **a**. Black, blue, and red ticks indicate spikes of neither, mouse-A, and mouse-B neurons, respectively. **d** Mouse-A neurons (blue) had significantly elevated peak firing rates during SPW-Rs compared with the neither neurons (black). The horizontal red line indicates the time period where the firing rate was significantly different (*p* < 0.01, permutation test). The shaded areas signify the SEMs. **e** Comparison of firing rates at ripple peaks (mean ± SEM). neither cells (N): *n* = 48 cells, 12.2 ± 1.6 Hz; mouse-A cells (A): *n* = 23 cells, 21.3 ± 3.6 Hz; \*\**p* = 0.0037, Student’s *t*-test. **f** The fraction of SPW-Rs for which a cell fired at least one spike, averaged across all cells. \*\**p* = 0.0048, Wilcoxon rank sum test. **g**, **h** Pairwise similarities of unit activities during SPW-Rs (**g**) and their statistical significance (**h**; permutation test). Note that units are sorted according to the social preferences plotted in **a**. **i**, **j** Comparison of the online (**i**) and offline (**j**) activity similarity between N/N (*n* = 92) and A/A (*n* = 23) cell pairs. \*\**p* = 0.017 and \*\*\**p* = 0.0045, Wilcoxon rank sum test. **k** Correlations between online and offline activity similarity. \*\**p* = 0.0058, permutation test (Supplementary Fig. 3). **l** Quantitative measurement and comparison of the offline similarities at different threshold MUA factors between N/N (*n* = 92) and A/A (*n* = 23) cell pairs. Main effect of Group, *F*_(1,103)_ = 19.4, *p* = 2.6×10^-5^, Group×Threshold, *F*_(1,103)_ = 17.0, *p* = 7.6×10^-5^, two-way repeated measures ANOVA. \*\*\**p* < 0.001, Tukey–Kramer multiple comparison test.

To measure the synchrony between cells during offline SPW-R events, we calculated the cosine similarity between pairs of spike count vectors. When the recorded units were sorted according to the online social preferences (Fig. 2g), the rendered square matrix of the similarity values clearly showed that a pair of familiar mouse-A cells (#1 and #2) were highly synchronized, to the exclusion of coactivation with other cells (Fig. 2g). In contrast, a pair of novel mouse-B cells (#6 and #7) were activated together with other cells during SPW-R events above the probability threshold calculated based on random shuffling (Fig. 2h), suggesting that the preferential reactivation of social memory neurons takes place at the population level. Likewise, by calculating the cosine similarity between spike count vector pairs in terms of video frames (25 Hz) during the online recordings, we found that pairs of mouse-A cells showed higher similarities than pairs of neither cells during both online (Fig. 2i) and offline recording (Fig. 2j). While pairs of both neither cells and familiar mouse-A cells showed significant positive correlations between online and offline similarities, the Pearson’s correlation coefficient of mouse-A cells was significantly higher than that of neither cells (*p* = 0.0058, permutation test) (Fig. 2c and Supplementary Fig. 3). Upon varying the threshold of the multi-unit activity (MUA) during SPW-Rs, we found that SPW-Rs with higher neuronal activities contained more correlated information content, and that this relationship was more prominent among mouse-A cells than neither cells (92 neither and 23 mouse-A cell pairs; main effect of Group, *F*_(1,103)_ = 19.4, *p* = 2.6×10^-5^; Group×Threshold, *F*_(1,103)_ = 17.0, *p* = 7.6×10^-5^; two-way repeated measures ANOVA) (Fig. 2l), again indicating that social memory was consolidated by the prioritized replay of specific neuronal ensembles.

### Firing sequence of social memory neurons

Dorsal hippocampus place cells encoding spatial representations are reactivated during SPW-Rs while maintaining the sequential firing order in which they were activated during previous spatial exploration [9–11]. To examine whether social memory neurons were analogously organized sequentially, we analyzed the temporal structure of vCA1 firing activities during offline SPW-R events. As a representative example, we recorded eight putative pyramidal cells and identified two of them as familiar mouse-A cells (Unit-#1 and 2) (Fig. 3a). When their offline similarities were sorted according to social preference scores, the square matrix showed that four cells, including the mouse-A cells, were synchronously activated during SPW-R events (Fig. 3b). By examining the temporal orders of the first spike of each unit observed during SPW-Rs, we observed that two sequences out of the 24 possible variations had a significantly higher probability of occurrence than that of a random event (Fig. 3c) and that the members of the spike sequence had respective preferred orders (Fig. 3d).

**Fig. 3.**
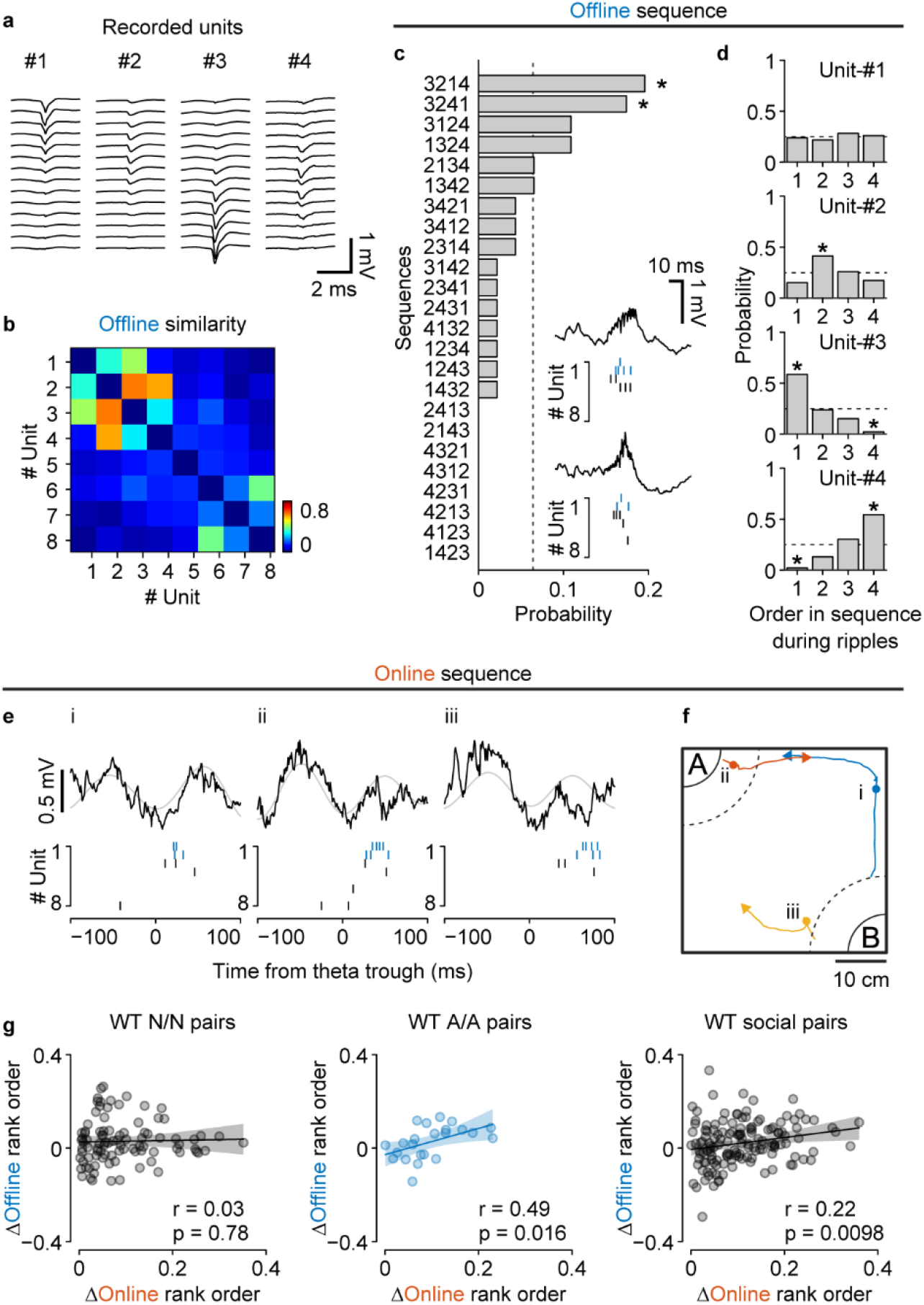
Spike sequences during online theta cycles and offline SPW-Rs. **a** Example four units recorded in the same session. **b** Activity similarity during offline SPW-Rs. Unit numbers correspond to those in **a**, and the units are sorted according to the social preferences. **c** Occurrence probability of the (4! = 24) possible neuronal sequences. The dotted line indicates chance level. **d** Probabilities that each unit was observed on a specific order within sequences. Dotted lines and gray bands indicate the chance levels. The asterisks in **c** and **d** represent motifs that are significantly more abundant or sparse than the chance (*p* < 0.05; determined by two-sided binomial test). **e** Example theta events during the SDT and simultaneously recorded spike sequences. Black and gray curves represent wide-band LFPs and theta (6-12 Hz) band signals, respectively. **f** Behavioral trajectories around the theta events shown in **e**. Circles depict the positions where the events occurred, and lines show behavioral paths of the subject 1.5 s before and after the events. Note that all example theta events occurred while either interacting with (i) or approaching (i, iii) the familiar social target, mouse-A. **g** Correlations between the differences of mean rank orders during online theta cycles and offline ripples. Orders within each cell pair were determined so that the Δonline mean rank orders always had a positive value. Pairs including at least one mouse-A or mouse-B cell were considered social pairs. Pairs of neither cells (N/N): *n* = 92 pairs, pairs of mouse-A cells (A/A): *n* = 23 pairs, social: *n* = 140 pairs.

To determine whether the spike sequences detected during offline SPW-Rs reflected the temporal structure of neuronal activities during online social experiences, we investigated the firing patterns of vCA1 neurons in terms of individual theta (6-12 Hz) cycles during the SDT, and found that similar activity patterns were preserved (Fig. 3e). Interestingly, these theta-locked sequences arose while the subject was approaching (Fig. 3e, i and iii) or interacting with (Fig. 3e, ii) the familiar social target, consistent with our observation that social memory neurons could respond to preferred social targets that are at a distance (Fig. 1). To compare the relationship between sequential ordering of neurons during individual online theta cycles and offline SPW-R events, we calculated the mean rank order during cycles/events [15] and the differences between each unit pair. We found that pairs of mouse-A cells, but not pairs of neither cells, showed a significant positive correlation between online and offline sequential ordering (Fig. 3g). The positive correlation still existed when pairs consisting of at least one social cell (namely one mouse-A or mouse-B cell) were incorporated. These results suggest that social representations in the vCA1 are temporally orchestrated at the population level centering around the social memory neurons, which maintain the sequential consistency from online social experiences to the offline memory consolidation phase.

### Impaired social memory and disrupted spike sequences in Shank3-KO mice

Social memory impairment is a hallmark of ASD both in human patients [27] and in mouse models [28]. To validate the reported impairment of social novelty preference in ASD model *Shank3*-KO mice [29], we conducted our SDT as described in Fig. 1a. As expected, the duration of approach of *Shank3*-KO mice for both the familiar mouse-A and the novel mouse-B was comparable (Fig. 4a,b), with a discrimination index not significantly different from zero (Fig. 4c). Consistent with these behavioral abnormalities, the proportion of identified mouse-A cells in Shank3-KO mice was significantly smaller than that in wild-type mice (*Shank3*-KO: mouse-A cells, 10.8 %; mouse-B cells, 10.8 %; neither cells: 78.4 %; *p* = 0.019, Z test of proportions) (Fig. 1h, 4d,e), although the firing rate of recorded pyramidal cells did not differ between wild-type and *Shank3*-KO mice (Fig. 4f). We also observed that SPW-Rs in *Shank3*-KO mice during offline recording (Fig. 4g) resembled those observed in wild-type mice (Fig. 4h) in terms of the ripple rate (wild-type: 0.15 ± 0.07 Hz, *n* = 11 sessions; *Shank3*-KO: 0.13 ± 0.07 Hz, *n* = 7 sessions; *p* = 0.33, Wilcoxon rank sum test), duration (wild-type: 53.5 ± 7.0 ms; *Shank3*-KO: 54.0 ± 5.1 ms; *t*^(16)^ = −0.15; *p* = 0.89, Student’s *t*-test), and peak frequency (wild-type: 155 ± 2.3 Hz, *n* = 11 sessions; *Shank3*-KO: 154 ± 1.7 Hz, *n* = 7 sessions; *t*_(16)_ = 0.25; *p* = 0.80, Student’s *t*-test) (Supplementary Fig. 4). Proportion of time spent resting (wild-type: 64 ± 3.4 %, *n* = 11 sessions; *Shank3*-KO: 75 ± 7.1%, *n* = 7 sessions; *t*_(16)_ = −1.6; *p* = 0.14, Student’s *t*-test) and the power spectrum density of the local field potential (LFP) signals during the offline recordings were also comparable (Supplementary Fig. 3). However, we found that the z-scored peak amplitude of SPW-R events was significantly lower in *Shank3*-KO mice compared to that in wild-type mice (wild-type: 6.8 ± 0.08, *n* = 11 sessions; *Shank3*-KO: 6.4 ± 0.09, *n* = 7 sessions; *t*_(16)_ = 3.17; *p* = 0.0056, Student’s *t*-test) (Fig. 4i,j). The difference was ascribed to the decrease in larger SPW-R events (Fig. 4k), indicating the absence of ripple power increase related to learning and memory consolidation[30].

**Fig. 4.**
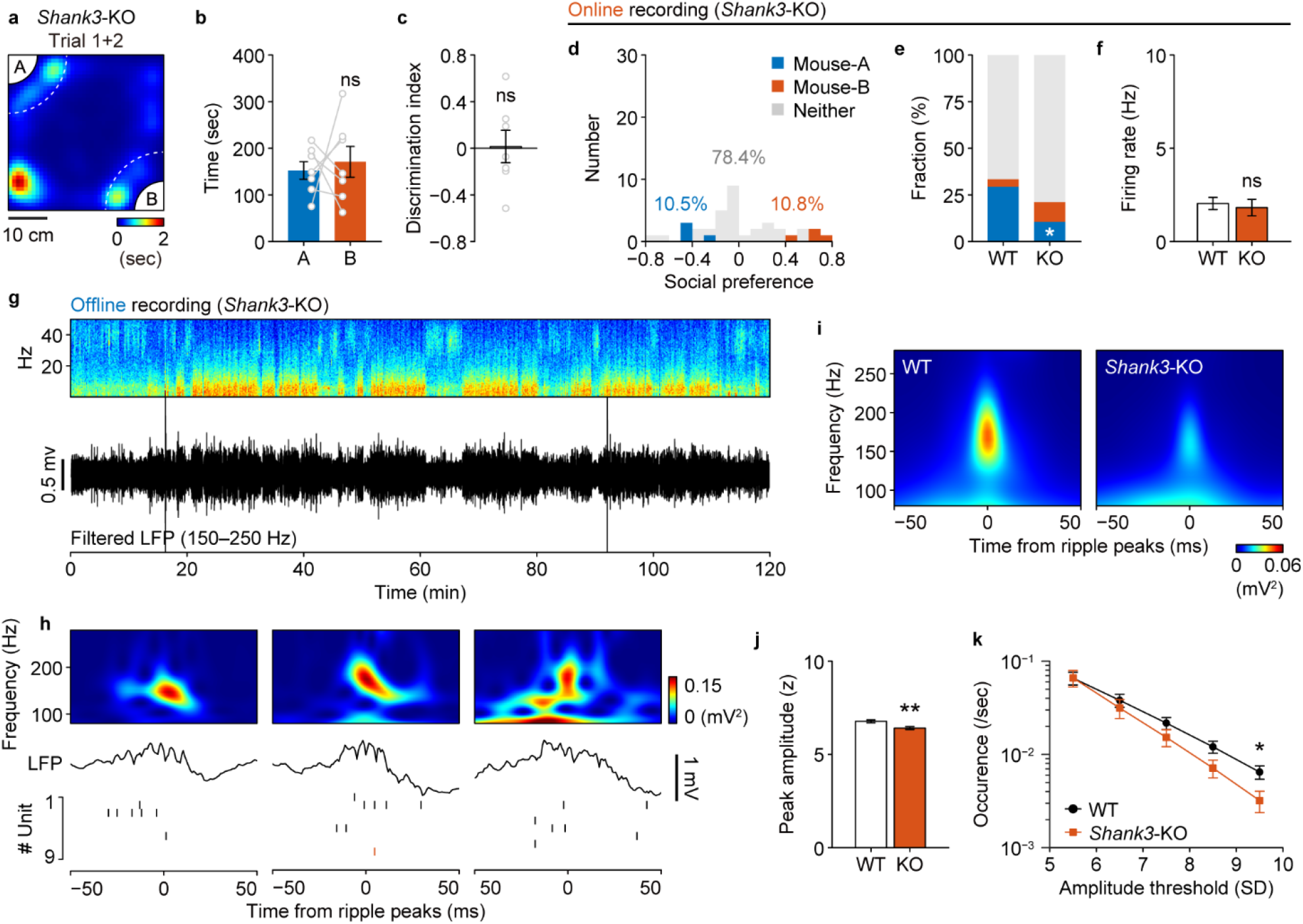
Impaired social memory and altered vCA1 activity of *Shank3*-KO mice. **a** Representative heat map of occupancy time during Trial 1 and Trial 2 with a *Shank3*-KO mouse. Note that the result of Trial 2 is rotated and superposed onto that of Trial 1. **b**, **c** Comparison of duration spent in the social zone (**b**) and the discrimination index (**c**). *n* = 7 with 4 subjects. Data are represented as mean ± SEM. **d** Social preference scores of recorded pyramidal neurons (*n* = 37 cells). Blue, mouse-A cell (*n* = 4 cells, 10.8 %); red, mouse-B cell (*n* = 4 cells, 10.8 %); grey neither neuron (*n* = 29 cells, 78.4 %). **e** Comparison of the fractions of neurons with significant social representation between WT and Shank3-KO mice. \**p* = 0.019, Z test of proportions. **f** Comparison of firing rates (mean ± SEM). **g** Example power spectrum and filtered ripple band trace of the LFP recorded in the home cage. **h** Representative SPW-Rs and raster plots of the units recorded during the same session as in **g**. Black and red ticks indicate spikes of neither and mouse-B neurons, respectively. **i** Representative SPW-R triggered spectrograms averaged across entire representative sessions from WT and KO mice. **j** Averaged peak ripple amplitude was significantly lower in KO mice than in WT mice. Data are represented as mean ± SEM. \*\**p* = 0.0056, Student’s *t-* test. **k** *Shank3*-KO mice generated fewer ripples with large amplitudes. \**p* < 0.05, Tukey–Kramer multiple comparison test.

Finally, we investigated the offline neuronal activity in the vCA1 of *Shank3*-KO mice and examined its consistency with the activity patterns during online social experiences (Fig. 5a,b). Contrary to our expectations, both online and offline similarities were indistinguishable from those of wild-type mice (Fig. 5c) and exhibited a significant positive correlation, comparable to that in the wild-type mice (Fig. 5d). However, by varying the threshold, we found that SPW-Rs with higher MUAs did not give rise to more correlated neuronal activities in *Shank3*-KO mice compared to that in wild-type mice (wild-type: *n* = 206 pairs, *Shank3*-KO: *n* = 100 pairs; main effect of Genotype, *F*_(1,304)_ = 9.7, *p* = 0.0020; Genotype×Threshold, *F*_(1,304)_ = 9.8, *p* = 0.0019) (Fig. 5e). Furthermore, the correlation between online and offline sequential ordering was disrupted in *Shank3*-KO mice (Fig. 5f). These results suggest that both populational and sequential consistency of neuronal activities between online social experiences and offline memory consolidation phases are impaired in *Shank3*-KO mice.

**Fig. 5.**
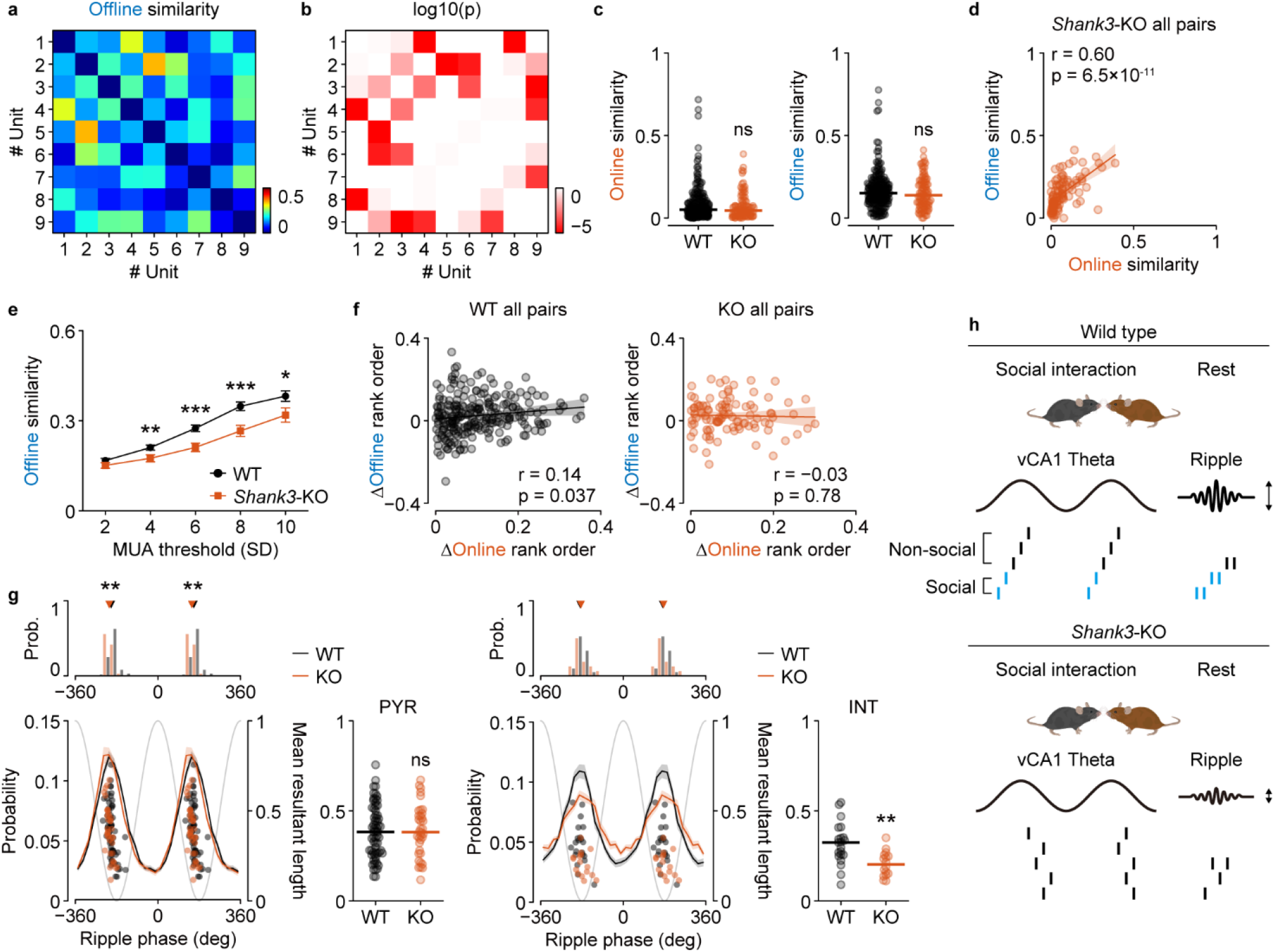
Disrupted spike sequences and ripple phase locking in *Shank3*-KO mice. **a**, **b** Pairwise similarities of unit activities during SPW-Rs (**a**) and their statistical significance (**b**; based on a permutation test). Units are sorted according to the social preferences. **c** Comparable online (light) and offline (right) activity similarity between pairs from WT (*n* = 237 pairs, 11 sessions) and KO (*n* = 100 pairs, 7 sessions) mice. **d** Correlations between online and offline activity similarity of simultaneously recorded cell pairs from KO mice. **e** Quantitative measurement and comparison of offline similarities at different MUA threshold factors between WT (*n* = 206 pairs) and KO (*n* = 100 pairs) mice. Main effect of Genotype, *F*_(1,304)_ = 9.7, *p* = 0.0020; Genotype×Threshold, *F*_(1,304)_ = 9.8, *p* = 0.0019. \*\*\**p* < 0.001, \*\**p* < 0.01, and \**p* < 0.05, Tukey–Kramer multiple comparison test. Data are represented as mean ± SEM. **f** Correlations between the differences of mean rank orders during online theta cycles and offline ripples. **g** Spike to ripple phase relationships of putative pyramidal cells (WT: *n* = 64 cells; KO: n = 36 cells) and interneurons (WT: *n* = 21 cells; KO: *n* = 16 cells). Note that only values from significantly (*p* < 0.001) phase-locked neurons were plotted. Shaded areas signify SEMs. Black and red triangles on the histograms represent median values of the preferred ripple phases among cells from wild-type and *Shank3*-KO mice, respectively. \*\**p* = 0.0035, circular multi-sample median test. Dot plots show the comparison of the strength of ripple phase preference. \*\**p* = 0.0017, Wilcoxon rank sum test. **h** Schematic representation of the findings.

Temporally precise, phase-locked inhibition supports hippocampal ripple generation [31, 32]. To investigate the neural substrate of disrupted sequential ordering, we compared the phase-locking of spike timings to ripple oscillations across groups. While the ripple oscillation induced a significant (*p* < 0.001, Rayleigh test) phase-locking in both the majority of pyramidal cells (wild-type: mean 156.3 ± 2.7°, 86.5% significantly phase-locked, *n* = 74 cells; Shank3-KO: mean 149.6 ± 2.1°, 97.3% significantly phase-locked, *n* = 37 cells; *p* = 0.08, Z test of proportions) and interneurons (wild-type: mean 176.7 ± 4.3°, 75.0% significantly phase-locked, *n* = 28 cells; *Shank3*-KO: mean 181.7 ± 8.6°, 80.0% significantly phase-locked, *n* = 20 cells; *p* = 0.68, Z test of proportions) across both genotypes, the strength of phase-locking (expressed as mean resultant length) was specifically decreased among interneurons (wild-type: 0.32 ± 0.03, *n* = 21 cells; *Shank3*-KO: 0.21 ± 0.02, *n* = 16 cells; *p* = 0.0017, Wilcoxon rank sum test) but not pyramidal cells (wild-type: 0.38 ± 0.03, *n* = 64 cells; *Shank3*-KO: 0.39 ± 0.02, *n* = 36 cells; *p* = 0.84, Wilcoxon rank sum test) in *Shank3*-KO mice (Fig. 5g). These results indicate that a loss of phase-locked inhibitory inputs onto pyramidal neurons may lead to an impairment of precise spike timings and their temporal sequences in social cell assemblies.

## DISCUSSION

Our findings demonstrated that vCA1 neuronal ensembles that encode memories of familiar social targets were preferentially reactivated by SPW-Rs that arise during the resting period after social interactions. A growing body of work examining social memory functions has focused primarily on two pivotal sub-regions of the hippocampus, the vCA1 and dorsal CA2 (dCA2) [3, 33, 34]. A recent study reported that dCA2 neurons respond to novel conspecifics during social interaction and are reactivated during SPW-Rs [35]. In contrast, our study revealed that the instantaneous emergence of neuronal activities associated with a novel individual during social exploration was not detected in vCA1 neurons, whereas robust online activation and later offline reactivation of neuronal ensembles were observed in response to familiarized individuals (Fig. 1h and Fig. 2). When encoding spatial representations, place-cell activity in the hippocampus stabilizes during spatial exploration [36], and recorded neurons in the dCA1 require a certain minimum level of experience to establish stable spatial representations [37]. Analogously, to form and store social memories, neural ensembles in the vCA1, but not those in the dCA2, may require social interaction of a sufficient duration to develop stable and observable neuronal activities that represent familiar individuals. Since CA2 neuronal projections innervate downstream hippocampal structures along the septotemporal axis [38], social information processed in the dCA2 appears to contribute to the consolidation of social memories in the vCA1 by propagating SPW-Rs [17]. As synapses on CA2 pyramidal neurons in adult mice are more resistant to the induction of long-term potentiation (LTP) compared to those on CA1 neurons [39], dCA2 and vCA1 neurons may cooperate while executing distinct functions, such as processing of social information in the dCA2 and encoding and storing of social memories in the vCA1.

As observed in previous studies, we confirmed that wild-type subject mice interacted longer with novel individuals than with familiar individuals (Fig. 1c,d). This memory-guided discriminatory social behavior is supported by projections from vCA1 social engram neurons to the nucleus accumbens (NAc) [5]. A recent study demonstrated that SPW-Rs occur asynchronously in the dorsal and ventral hippocampus during the awake state and modulate distinct populations of NAc medium spiny neurons (MSNs) [40]. It has also been reported that dorsal but not ventral hippocampal SPW-Rs activate NAc neurons that encode information related to spatial and reward-related information [40]; therefore, ventral hippocampal SPW-Rs may specifically convey social information to MSNs in the NAc to establish a neural circuit that supports memory-guided social behavior.

Sequential activation of neuronal ensembles appears to serve as a general mechanism for the consolidation of semantic memories in the hippocampus [41]. In the dorsal hippocampus, firing sequences of neuronal ensembles during SPW-Rs reflect the continuous spatial trajectories of recent experiences [9–11], which can initially be measured as the successive activation of place cells at the nested time scales of behavior and the theta cycle [42]. In this study, we demonstrated that, in contrast with the non-social neurons, pairs of vCA1 social memory neurons activated in response to a familiar individual exhibited higher levels of coactivation during both online social interactions and offline SPW-Rs. Furthermore, spike sequences of vCA1 social memory neurons within online theta cycles were preserved during offline SPW-Rs, which indicated that social information likely employs temporal coding. Analogous to the place cell sequences in the dCA1, the sequential firing of vCA1 social memory neurons could reflect a certain type of information structure. One possible hypothesis is that the firing sequence might be organized according to the perceptual origin of social stimuli. Both in rodents and humans [43], social recognition and its memorization are supported by multimodal sensory inputs such as visual [44] and olfactory [45] stimuli. Indeed, “concept cells” in the human hippocampus that encode social memory can fire selectively not only in response to pictures of a specific person but also in response to other sensory modality cues such as the pronounced or written name of that person [46]. The presence of social neurons that are preferentially activated in response to specific social targets, even when the test subjects were distant from the social interaction zone (Fig. 1j), indicates that some vCA1 social memory neurons are more specifically tuned to visual or auditory cues, which can be processed across distances, rather than olfactory cues, which require close proximity. Nevertheless, future studies are necessary to examine whether and how social cues that are represented through a single sensory modality activate vCA1 social neurons, presumably through the dCA2-vCA1 pathway [38] or the conventional vCA3-vCA1 pathway, in which pattern separation and completion are thought to occur [47].

To consolidate an episodic memory, the offline reactivation of a neuronal ensemble must incorporate multiple features of the experience, including spatial, temporal, and social information. In line with this notion, our recordings captured sequential activities among both social and non-social neurons (Fig. 3). Neuronal recordings from a larger neuronal population could elucidate how the detailed contents of experiences are mapped onto sequential neural activities.

Impairments in social reciprocity result in the debilitating symptoms associated with several neurodevelopmental disorders, including ASD [48, 49]. *Shank3*, one of the most promising ASD-susceptibility genes [50], encodes a postsynaptic scaffolding protein [51], and ASD model mice generated by deleting *Shank3* are characterized by reduced social interactions and abnormal social novelty recognition [29, 52]. As in a human study which reported that adult patients with ASD present with deficiencies in their memory of faces and social scenes [27], we observed that homozygous *Shank3*-KO mice exhibited social memory impairments within our social discrimination paradigm (Fig. 4b,c). Consistent with this behavioral phenotype, a smaller proportion of vCA1 neurons in *Shank3*-KO mice responded to familiar individuals, even after forming social memory (Fig. 4d,e), although the overall firing rate of recorded pyramidal neurons was normal. How does the deficiency in Shank3 expression lead to these behavioral and physiological impairments? The induction of synaptic potentiation is thought to be a general mechanism underlying memory formation [53]. Several in vitro electrophysiological studies have reported that LTP is impaired in the hippocampal CA1 synapses of *Shank3*-KO mice, whereas long-term depression (LTD) remains unaffected [54]. Therefore, the synapses on social memory neurons in the vCA1 are likely to be similarly defective in terms of the induction of the necessary synaptic potentiation required to form a social memory.

More importantly, in our study, which focused on the neural coding that occurs during SPW-Rs, pairwise measures of unit coactivation revealed that *Shank3*-KO mice showed decreased coactivation of vCA1 neurons during SPW-Rs, likely due to the malformation of neural networks resulting from the impairment of LTP on vCA1 neurons. A series of studies have demonstrated that the appropriate balance between the LTP and LTD of CA1 neurons induced by SPW-Rs is crucial for memory formation and consolidation. The relative spike timings of dCA3 and dCA1 place cells during SPW-Rs induce synaptic potentiation on dCA1 neurons [55]. In contrast, SPW-Rs that occur during slow-wave sleep lead to spontaneous synaptic depression on dCA1 neurons, which is dependent on the N-methyl-D-aspartate receptor, whose inactivation results in impaired learning of new memories [14]. Schizophrenia model Calcineurin-KO mice, which are severely LTD-deficient at hippocampal synapses, showed impaired ripple-associated reactivation of spatial sequences [56] and deficits in hippocampus-dependent working and episodic-like memory tasks [57]. Our results using an ASD model mouse indicated that malformations in vCA1 neural ensembles are likely due to the reduced tuning of SPW-Rs, which interferes with the formation of social memories and the elicitation of social familiarity behaviors in patients with ASD.

In conclusion, our study provides empirical evidence to suggest that a key concept of social memory representation involves neural ensembles in the vCA1 and that pathological involvement of the vCA1 may underlie the physiological characteristics of ASD.

## DATA AVAILABILITY

The data supporting the findings in the present study are available from the corresponding author upon reasonable request.

## ACKNOWLEDGEMENTS

We thank I. Yoshimura, A. Watarai, T.M. Tang, and M. Watanabe for technical assistance; and all members of the Okuyama laboratory for their support. We thank J. Yamamoto and D.S. Roy for their feedback on the manuscript. This work was supported by JST, PRESTO Grant Number JPMJPR1781, Japan (to TO); JSPS KAKENHI Grant Numbers 18H02544 (to TO) and 19K06951 (to KT); Takeda Science Foundation (to TO); The Uehara Memorial Foundation (to TO); The NOVARTIS Foundation (Japan) for the Promotion of Science (to TO); Daiichi Sankyo Foundation of Life Science (to TO); The Mochida Memorial Foundation for Medical and Pharmaceutical Research (to TO); The Naito Foundation (to TO); SECOM Science and Technology Foundation (to TO).

## AUTHOR CONTRIBUTIONS

KT and TO conceptualized and designed the study. KT, AW, and MC performed the experiments. KT and ZH analyzed the data. KT, MW, and TO wrote the manuscript. KT and TO contributed to funding acquisition. All the authors discussed the results and contributed to the manuscript.

## ADDITIONAL INFORMATION

### Supplementary Information

**Supplementary Fig. 1** Properties of vCA1 social cells.

**Supplementary Fig. 2** Sleep analysis and ripple event detection.

**Supplementary Fig. 3** A permutation test for comparing correlation coefficients.

**Supplementary Fig. 4** Properties of vCA1 neurons in *Shank3*-KO mice.

## SUPPLEMENTARY INFORMATION

**Supplementary Fig. 1.**
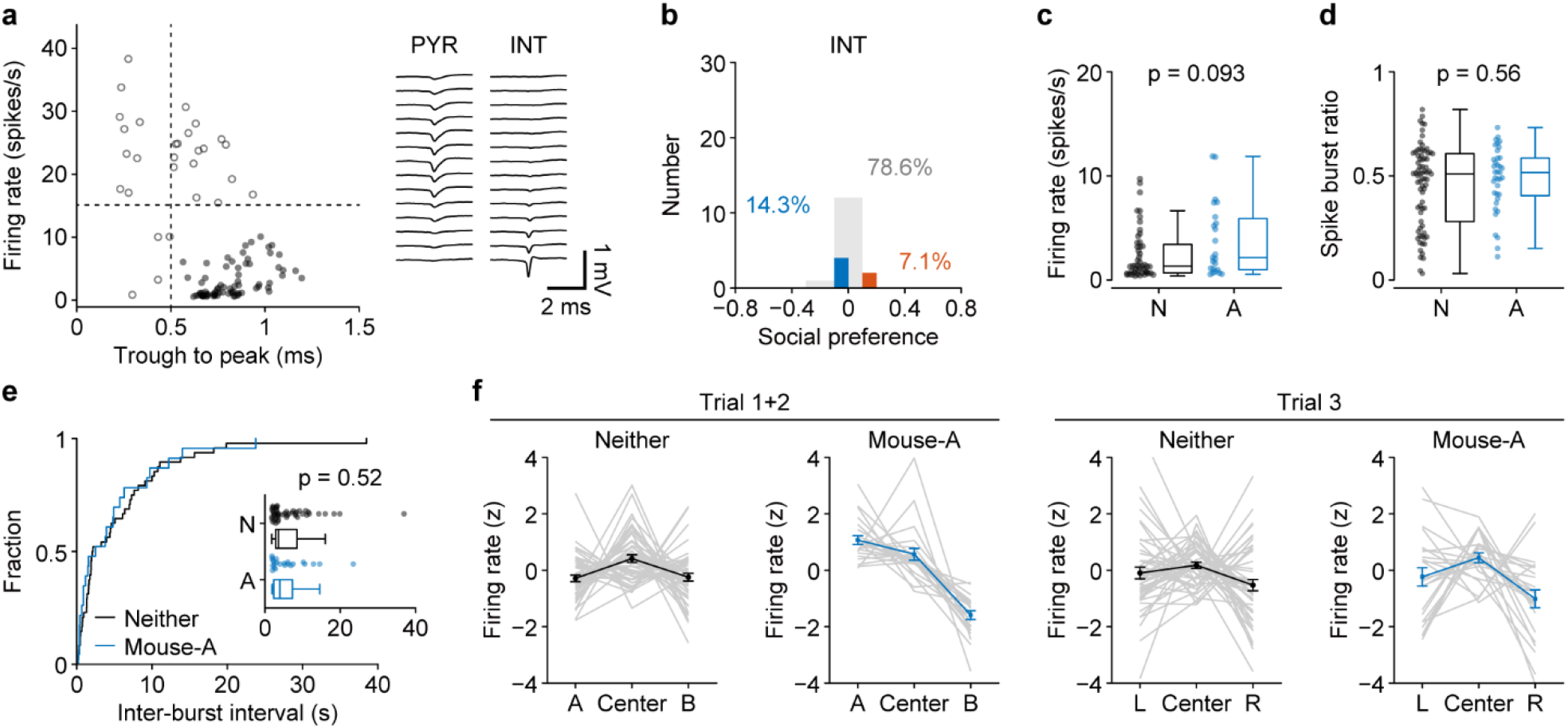
Properties of vCA1 social cells. **a** Neurons with a low firing rate (< 15 Hz) and wide trough-to-peak spike shape length (> 0.5 ms) were classified as excitatory pyramidal neurons. **b** Social preference scores of recorded interneurons (*n* = 29 cells). **c** Firing rate of neither cells (N; *n* = 48) and mouse-A cells (A; *n* = 23). **d** Proportion of spikes in bursts (spikes occurring with an inter-spike interval of 3–15 ms) of neither cells and mouse-A cells. **e** Distribution of inter-spike-intervals of neither cells and mouse-A cells. **f** Z-scored firing rates of mouse-A cells and neither cells during the social trials (Trial 1+2; left) and the control trial (Trial 3; right).

**Supplementary Fig. 2.**
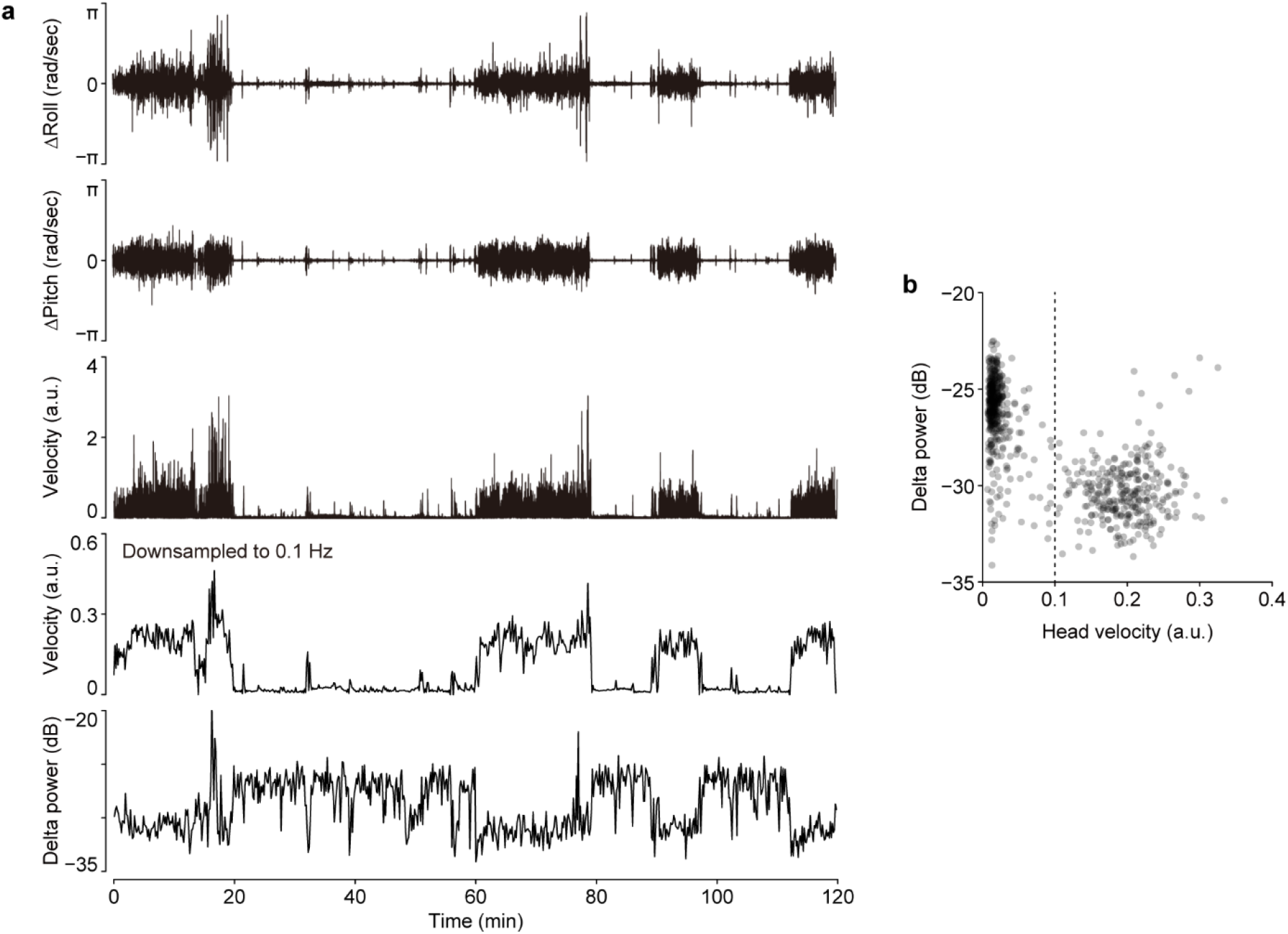
Sleep analysis and ripple event detection. **a** Representative signals related to head movement. Signals from a 3-axis accelerometer mounted on the headstage were used to calculate the derivatives of roll and pitch of the subject’s head, and the L2 norm of these derivatives was used as a proxy for the overall movement of the subject. **b** Relationship between head movement and the instantaneous delta power recorded from the vCA1. Bin length = 10 s. Epochs with head velocity less than 0.1 arbitrary unit (a.u.) were considered to be immobile, sleeping periods.

**Supplementary Fig. 3.**
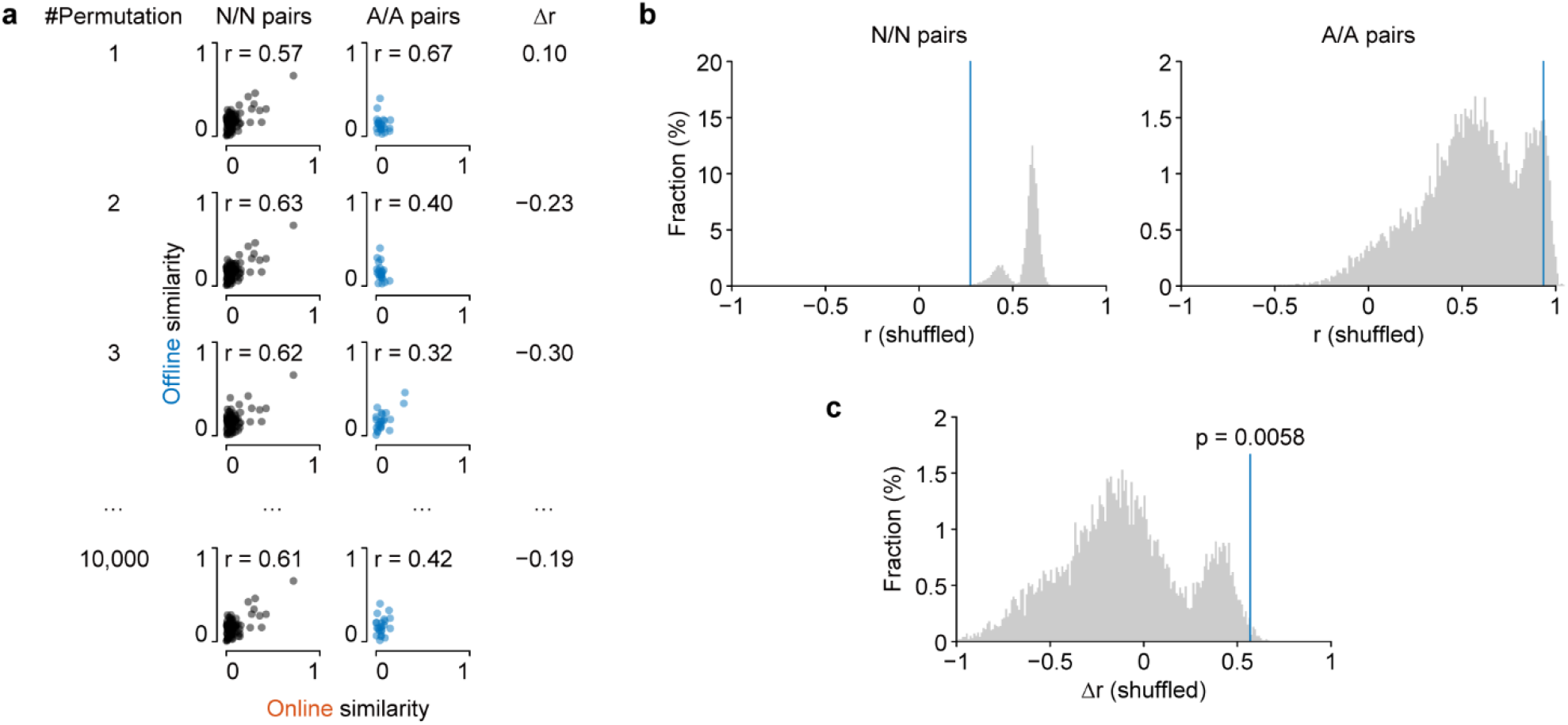
A permutation test for comparing correlation coefficients. **a** Pairs of neither cells (*n* = 92 pairs) and mouse-A cells (*n* = 23 pairs) were shuffled and randomly divided into two groups with the same number of pairs with the actual data. Correlation coefficients for each shuffled group and the difference between them were calculated, and this procedure was repeated 10,000 times. **b**,**c** Frequency distributions of the coefficients calculated for the shuffled groups were compared with the original values to test two-tailed significance.

**Supplementary Fig. 4.**
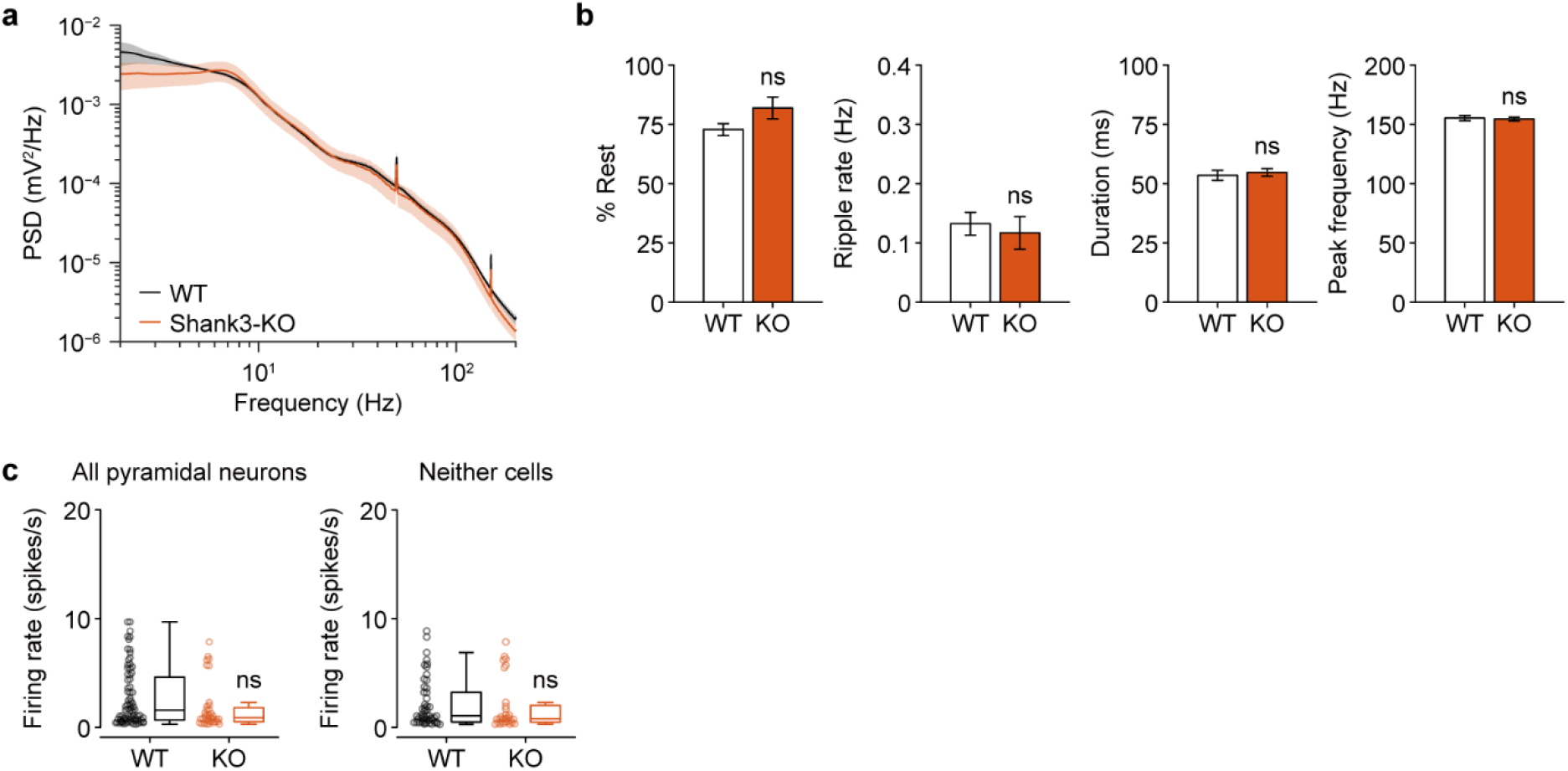
Properties of vCA1 neurons in Shank3-KO mice. **a** Power spectrum density of the LFP signals recorded from the vCA1 of WT (*n* = 11 sessions) and KO (*n* = 7 sessions) mice. **b** Proportion of time spent resting, ripple rate, ripple duration, and peak ripple frequency were comparable between WT and KO mice. **c** Firing rate of vCA1 neurons during the SDT was comparable between WT (*n* = 74 cells) and KO (*n* = 37 cells) mice.

## REFERENCES

1. Walum H, Young LJ. The neural mechanisms and circuitry of the pair bond. Nat Rev Neurosci. 2018;19:643–654.

2. Okuyama T, Yokoi S, Abe H, Isoe Y, Suehiro Y, Imada H, et al. A neural mechanism underlying mating preferences for familiar individuals in medaka fish. Science. 2014;343:91–94.

3. Hitti FL, Siegelbaum SA. The hippocampal CA2 region is essential for social memory. Nature. 2014;508:88–92.

4. Remedios R, Kennedy A, Zelikowsky M, Grewe BF, Schnitzer MJ, Anderson DJ. Social behaviour shapes hypothalamic neural ensemble representations of conspecific sex. Nature. 2017;550:388–392.

5. Okuyama T, Kitamura T, Roy DS, Itohara S, Tonegawa S. Ventral CA1 neurons store social memory. Science. 2016;353:1536–1541.

6. Rao RP, von Heimendahl M, Bahr V, Brecht M. Neuronal Responses to Conspecifics in the Ventral CA1. Cell Rep. 2019;27:3460–3472 e3.

7. Okuyama T. Social memory engram in the hippocampus. Neurosci Res. 2018;129:17–23.

8. Buzsaki G. Hippocampal sharp wave-ripple: A cognitive biomarker for episodic memory and planning. Hippocampus. 2015;25:1073–1188.

9. Wilson MA, McNaughton BL. Reactivation of hippocampal ensemble memories during sleep. Science. 1994;265:676–679.

10. Nádasdy Z, Hirase H, Czurkó A, Csicsvari J, Buzsáki G. Replay and time compression of recurring spike sequences in the hippocampus. J Neurosci. 1999;19:9497–9507.

11. Lee AK, Wilson MA. Memory of sequential experience in the hippocampus during slow wave sleep. Neuron. 2002;36:1183–1194.

12. O’Neill J, Senior T, Csicsvari J. Place-selective firing of CA1 pyramidal cells during sharp wave/ripple network patterns in exploratory behavior. Neuron. 2006;49:143–155.

13. Foster DJ, Wilson MA. Reverse replay of behavioural sequences in hippocampal place cells during the awake state. Nature. 2006;440:680–683.

14. Norimoto H, Makino K, Gao M, Shikano Y, Okamoto K, Ishikawa T, et al. Hippocampal ripples down-regulate synapses. Science. 2018;359:1524–1527.

15. Fernández-Ruiz A, Oliva A, Fermino de Oliveira E, Rocha-Almeida F, Tingley D, Buzsáki G. Long-duration hippocampal sharp wave ripples improve memory. Science. 2019;364:1082–1086.

16. Girardeau G, Benchenane K, Wiener SI, Buzsáki G, Zugaro MB. Selective suppression of hippocampal ripples impairs spatial memory. Nat Neurosci. 2009;12:1222–1223.

17. Patel J, Schomburg EW, Berényi A, Fujisawa S, Buzsáki G. Local generation and propagation of ripples along the septotemporal axis of the hippocampus. J Neurosci. 2013;33:17029–17041.

18. Ciocchi S, Passecker J, Malagon-Vina H, Mikus N, Klausberger T. Brain computation. Selective information routing by ventral hippocampal CA1 projection neurons. Science. 2015;348:560–563.

19. Patel J, Fujisawa S, Berényi A, Royer S, Buzsáki G. Traveling theta waves along the entire septotemporal axis of the hippocampus. Neuron. 2012;75:410–417.

20. Lopes G, Bonacchi N, Frazão J, Neto JP, Atallah BV, Soares S, et al. Bonsai: an event-based framework for processing and controlling data streams. Front Neuroinform. 2015;9:7.

21. Mathis A, Mamidanna P, Cury KM, Abe T, Murthy VN, Mathis MW, et al. DeepLabCut: markerless pose estimation of user-defined body parts with deep learning. Nat Neurosci. 2018;21:1281–1289.

22. Siegle JH, López AC, Patel YA, Abramov K, Ohayon S, Voigts J. Open Ephys: an open-source, plugin-based platform for multichannel electrophysiology. J Neural Eng. 2017;14:045003.

23. Pachitariu M, Steinmetz NA, Kadir SN, Carandini M, Harris KD. Fast and accurate spike sorting of high-channel count probes with KiloSort. In: Lee D, Sugiyama M, Luxburg U, Guyon I, Garnett R, editors. Advances in Neural Information Processing Systems, vol. 29, Curran Associates, Inc.; 2016. p. 4448–4456.

24. Rossant C, Kadir SN, Goodman DFM, Schulman J, Hunter MLD, Saleem AB, et al. Spike sorting for large, dense electrode arrays. Nat Neurosci. 2016;19:634–641.

25. Barthó P, Hirase H, Monconduit L, Zugaro M, Harris KD, Buzsáki G. Characterization of neocortical principal cells and interneurons by network interactions and extracellular features. J Neurophysiol. 2004;92:600–608.

26. Stark E, Eichler R, Roux L, Fujisawa S, Rotstein HG, Buzsáki G. Inhibition-induced theta resonance in cortical circuits. Neuron. 2013;80:1263–1276.

27. Williams DL, Goldstein G, Minshew NJ. Impaired memory for faces and social scenes in autism: clinical implications of memory dysfunction. Arch Clin Neuropsychol. 2005;20:1–15.

28. Ferguson JN, Young LJ, Hearn EF, Matzuk MM, Insel TR, Winslow JT. Social amnesia in mice lacking the oxytocin gene. Nat Genet. 2000;25:284–288.

29. Peça J, Feliciano C, Ting JT, Wang W, Wells MF, Venkatraman TN, et al. Shank3 mutant mice display autistic-like behaviours and striatal dysfunction. Nature. 2011;472:437–442.

30. Eschenko O, Ramadan W, Mölle M, Born J, Sara SJ. Sustained increase in hippocampal sharp-wave ripple activity during slow-wave sleep after learning. Learn Mem. 2008;15:222–228.

31. Stark E, Roux L, Eichler R, Senzai Y, Royer S, Buzsáki G. Pyramidal cell-interneuron interactions underlie hippocampal ripple oscillations. Neuron. 2014;83:467–480.

32. Gan J, Weng S-M, Pernía-Andrade AJ, Csicsvari J, Jonas P. Phase-Locked Inhibition, but Not Excitation, Underlies Hippocampal Ripple Oscillations in Awake Mice In Vivo. Neuron. 2017;93:308–314.

33. Watarai A, Tao K, Wang M-Y, Okuyama T. Distinct functions of ventral CA1 and dorsal CA2 in social memory. Curr Opin Neurobiol. 2021;68:29–35.

34. Smith AS, Williams Avram SK, Cymerblit-Sabba A, Song J, Young WS. Targeted activation of the hippocampal CA2 area strongly enhances social memory. Mol Psychiatry. 2016;21:1137–1144.

35. Oliva A, Fernandez-Ruiz A, Leroy F, Siegelbaum SA. Hippocampal CA2 sharp-wave ripples reactivate and promote social memory. Nature. 2020;2020/09/25.

36. Wilson MA, McNaughton BL. Dynamics of the hippocampal ensemble code for space. Science. 1993;261:1055–1058.

37. Frank LM, Stanley GB, Brown EN. Hippocampal plasticity across multiple days of exposure to novel environments. J Neurosci. 2004;24:7681–7689.

38. Meira T, Leroy F, Buss EW, Oliva A, Park J, Siegelbaum SA. A hippocampal circuit linking dorsal CA2 to ventral CA1 critical for social memory dynamics. Nat Commun. 2018;9:4163.

39. Carstens KE, Dudek SM. Regulation of synaptic plasticity in hippocampal area CA2. Curr Opin Neurobiol. 2019;54:194–199.

40. Sosa M, Joo HR, Frank LM. Dorsal and Ventral Hippocampal Sharp-Wave Ripples Activate Distinct Nucleus Accumbens Networks. Neuron. 2020;105:725–741.e8.

41. Dragoi G. Cell assemblies, sequences and temporal coding in the hippocampus. Curr Opin Neurobiol. 2020;64:111–118.

42. Drieu C, Todorova R, Zugaro M. Nested sequences of hippocampal assemblies during behavior support subsequent sleep replay. Science. 2018;362:675–679.

43. Chung M, Wang M-Y, Huang Z, Okuyama T. Diverse sensory cues for individual recognition. Dev Growth Differ. 2020. 28 October 2020. https://doi.org/10.1111/dgd.12697.

44. Watanabe S, Shinozuka K, Kikusui T. Preference for and discrimination of videos of conspecific social behavior in mice. Anim Cogn. 2016;19:523–531.

45. Spehr M, Kelliher KR, Li X-H, Boehm T, Leinders-Zufall T, Zufall F. Essential role of the main olfactory system in social recognition of major histocompatibility complex peptide ligands. J Neurosci. 2006;26:1961–1970.

46. Quiroga RQ. Concept cells: the building blocks of declarative memory functions. Nat Rev Neurosci. 2012;13:587–597.

47. Rolls ET. An attractor network in the hippocampus: theory and neurophysiology. Learn Mem. 2007;14:714–731.

48. Geschwind DH, Levitt P. Autism spectrum disorders: developmental disconnection syndromes. Curr Opin Neurobiol. 2007;17:103–111.

49. Wing L, Gould J. Severe impairments of social interaction and associated abnormalities in children: epidemiology and classification. J Autism Dev Disord. 1979;9:11–29.

50. Durand CM, Betancur C, Boeckers TM, Bockmann J, Chaste P, Fauchereau F, et al. Mutations in the gene encoding the synaptic scaffolding protein SHANK3 are associated with autism spectrum disorders. Nat Genet. 2007;39:25–27.

51. Naisbitt S, Kim E, Tu JC, Xiao B, Sala C, Valtschanoff J, et al. Shank, a novel family of postsynaptic density proteins that binds to the NMDA receptor/PSD-95/GKAP complex and cortactin. Neuron. 1999;23:569–582.

52. Qin L, Ma K, Wang Z-J, Hu Z, Matas E, Wei J, et al. Social deficits in Shank3-deficient mouse models of autism are rescued by histone deacetylase (HDAC) inhibition. Nat Neurosci. 2018;21:564–575.

53. Whitlock JR, Heynen AJ, Shuler MG, Bear MF. Learning induces long-term potentiation in the hippocampus. Science. 2006;313:1093–1097.

54. Bozdagi O, Sakurai T, Papapetrou D, Wang X, Dickstein DL, Takahashi N, et al. Haploinsufficiency of the autism-associated Shank3 gene leads to deficits in synaptic function, social interaction, and social communication. Mol Autism. 2010;1:15.

55. Sadowski JHLP, Jones MW, Mellor JR. Sharp-Wave Ripples Orchestrate the Induction of Synaptic Plasticity during Reactivation of Place Cell Firing Patterns in the Hippocampus. Cell Rep. 2016;14:1916–1929.

56. Suh J, Foster DJ, Davoudi H, Wilson MA, Tonegawa S. Impaired hippocampal ripple-associated replay in a mouse model of schizophrenia. Neuron. 2013;80:484–493.

57. Zeng H, Chattarji S, Barbarosie M, Rondi-Reig L, Philpot BD, Miyakawa T, et al. Forebrain-specific calcineurin knockout selectively impairs bidirectional synaptic plasticity and working/episodic-like memory. Cell. 2001;107:617–629.

